# The dynamic modular fingerprints of the human brain at rest

**DOI:** 10.1101/2020.05.30.125385

**Authors:** Aya Kabbara, Veronique Paban, Mahmoud Hassan

**Author notes:** Corresponding author: Mahmoud Hassan.

## Abstract

The human brain is a dynamic modular network that can be decomposed into a set of modules and its activity changes permanently over time. At rest, several brain networks, known as Resting-State Networks (RSNs), emerge and cross-communicate even at sub-second temporal scale. Here, we seek to decipher the fast reshaping in spontaneous brain modularity and its relationship to RSNs. We use Electro/Magneto-Encephalography (EEG/MEG) to track dynamics of modular brain networks, in three independent datasets (N= 568) of healthy subjects at rest. We show the presence of striking spatiotemporal network pattern consistent over participants. We also show that some RSNs, such as default mode network and temporal network, are not necessary ‘unified units’ but rather can be divided into multiple sub-networks over time. Using the resting state questionnaire, our results revealed also that brain network dynamics are strongly correlated to mental imagery at rest. These findings add new perspectives to brain dynamic analysis and highlight the importance of tracking fast reconfiguration of electrophysiological networks at rest.

## Introduction

Spontaneous brain activity changes permanently, over multiple temporal scales ranging from sub-second to years. This happens across a set of networks known as resting state networks (RSNs) (Damoiseaux et al., 2012; De Luca et al., 2005; Fox and Raichle, 2007; Greicius et al., 2003; Raichle et al., 2001). To decipher the ultra-fast dynamic reconfiguration of these RSNs and their cross-communications, several functional studies have been conducted. Some studies have described the dynamic topological changes of functional networks using graph theoretical analysis (Jiao et al., 2018; Kabbara et al., 2017a; Pasquale et al., 2010). Others focused on detecting ‘brain network states’ fluctuating over time. The expanding idea is that spontaneous brain activity can be explained by a set of spatiotemporal network pattern. This was often performed in combination with dimensionality reduction algorithms (such as K-means clustering (Allen et al., 2014; Damaraju et al., 2014), principal component analysis (Preti and Van De Ville, 2016) or orthogonal connectivity factorization (Hyvärinen et al., 2016)), or model-based approaches, such as hidden markov model (Baker et al., 2014; Vidaurre et al., 2017). Features derived from these fast-dynamic analyses were also shown as potential markers for some brain diseases (Filippi et al., 2019; A Kabbara et al., 2018; Liu et al., 2019) and behavioral characteristics (Kenett et al., 2020; Tompson et al., 2018).

Emerging evidence show that the human brain is a modular network, that is, it can be decomposed into a set of ‘modules’ (also called communities or clusters) denoting brain regions highly intra-connected and weakly connected with others (Bassett and Sporns, 2017; Bullmore and Sporns, 2009; Sporns and Betzel, 2016). The modular organization of the brain network and its dynamics was shown to be associated with aging (Meunier et al., 2009) and several task-related brain functions such as learning (Bassett et al., 2011) and cognitive effort (Kitzbichler et al., 2011).While several resting-state functional magnetic resonance imaging (fMRI) studies have been established to investigate the time-dependence of brain modular networks (Allen et al., 2014; Jones et al., 2012; Zalesky et al., 2014), the evidence for rapid reshaping in spontaneous modular brain networks and their relationships with RSNs at timescales associated with fast cognition is very limited. To effectively track network dynamics, we need a modality which can match the rapid timescales of the brain. Thus, it is mandatory to use electro/magneto-encephalography (EEG/MEG), which offer sub-millisecond temporal resolution, in order to describe such fast modularity-dependent fluctuations.

Here, we hypothesized that the dynamic modular reorganization of the human brain at rest is characterized by a continuous process of separation and merging within and across different RSNs over time. To detect “modular brain states”, we use a recently developed framework that allow to precisely quantify these time-varying ‘states’ fluctuations (A. Kabbara et al., 2019). Unlike other clustering algorithms, this framework looks at the fast-transient changes in the brain modular structure and was shown to outperform other existing clustering algorithms in terms of spatiotemporal precision ^26^. We tested our hypothesis on three independent EEG/MEG datasets (N=568) for healthy subjects at rest, source-reconstructed to 68 regions across the entire cortex. Dynamic brain networks were reconstructed using the EEG/MEG source connectivity technique using both power and phase-couplings(Mahmoud Hassan and Wendling, 2018a), combined with a sliding window approach and an algorithm to detect modular states (Figure 1). Notably, results revealed the presence of very consistent network patterns for most participants and a splitting phenomenon of some of these networks, such as the default mode network (DMN) and temporal network. We consider that tracking fast modular architecture of ongoing neuronal activity provides new insights into the dynamics of large-scale electrophysiological network organization of the brain.

**Figure 1.**
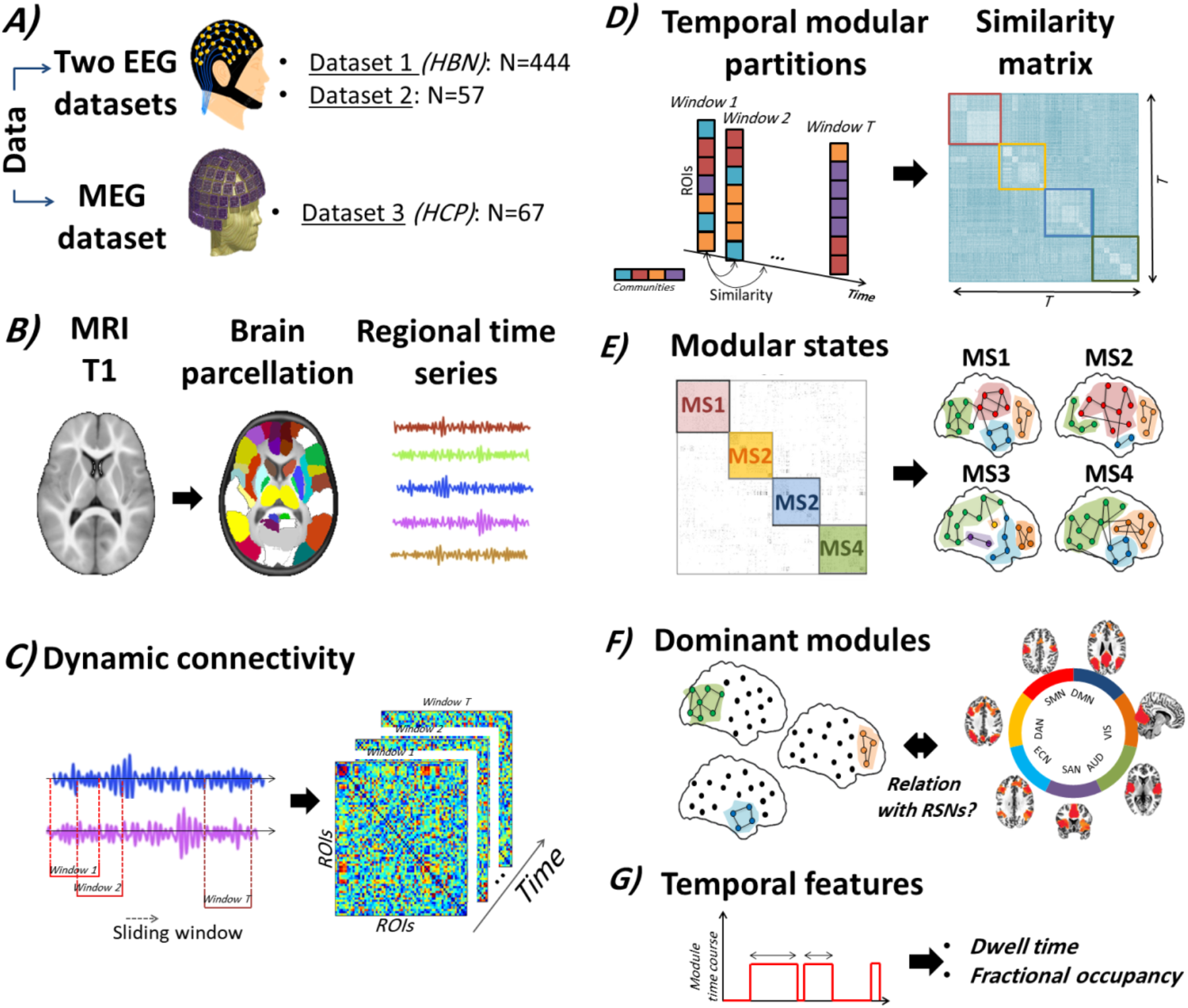
The full pipeline of the study. (A) Data analyzed comprised three datasets: 1) Resting state EEG data provided from the Healthy brain network biobank comprising 444 subjects, 2) Resting state EEG data acquired from 56 subjects and Resting state MEG data provided from the Human Connectome Project including 61 subjects. (B) The template MRI image was segmented into regions of interest (ROIs) by the means of an anatomical atlas (Desikan et al., 2006). Then, the regional time series of each subject were reconstructed using the weighted minimum norm estimate inverse solution (WMNE) for Datasets 1 and 2, and beamforming for Dataset 3. (C) Using a sliding window technique, the dynamic brain networks were computed. (D) The first step in the modularity-based algorithm is to parcellate each temporal network into communities. Then, the similarity between different temporal modular structures is assessed. (E) The similarity matrix is segmented into different communities where each one represents a modular state of specific spatial topology combining different time windows. (F) Following this, the dominant modules (common modules between different MSs) are extracted for each subject. The correspondence between these modules and the well-known RSNs is also detected. (G) The mean dwell time and fractional occupancies are calculated for the derived dominant modules.

## Materials and methods

### 1. EEG datasets

#### 1.1. Dataset 1 (HBN)

##### 1.1.1. Participants

As part of the Healthy Brain Network (HBN) Biobank release 1 (Alexander et al., 2017) http://fcon_1000.projects.nitrc.org/indi/cmi_healthy_brain_network/sharing_neuro.html, resting-state EEG data were collected from 444 healthy subjects (239 female). The release originally included 603 subjects, but data from 159 subjects were rejected after pre-processing and visual inspection. Subjects are healthy and aged between 5 and 21 years old. The Ids of the 444 participants are listed in Table S5 (see Supplementary materials).

##### 1.1.2. Data acquisition and pre-processing

High-density EEG data are recorded in a sound-shielded room at a sampling rate of 500 Hz with a bandpass of 0.1 to 100 Hz, using a 128-channel EEG geodesic hydrocel system by EGI. The recording reference is at Cz (vertex of the head). The impedance of each electrode is checked prior to recording, to ensure good contact, and is kept below 40 kOhm. Each EEG session consisted of 5 min resting period. As provided by the HBN, EEG signals were preprocessed using Automagic Matlab toolbox(Pedroni et al., 2019), visual inspection was also done on the data after automatic preprocessing. Briefly, it consists of interpolating the noisy, flat or outlier channels. The Multiple Artifact Rejection Algorithm (MARA) which automatizes the process of independent component analysis (ICA) was used to detect and reject artifacts such as the eye blinks and the movement artifacts (Winkler et al., 2011). Then, four artifact-free epochs of 40-s length were selected for each participant. This epoch length was used in a previous study, and was considered as a good compromise between the needed temporal resolution and the results reproducibility (Kabbara et al., 2017b).

#### 1.2. Dataset 2

##### 1.2.1. Participants

A total of 56 healthy subjects were recruited (29 female). The mean age was 34.7 years old (*SD* = 9.1 years, range = 18–55). Education ranged from 10 years of schooling to a PhD degree. None of the volunteers reported taking any medication or drugs, nor suffered from any past or present neurological or psychiatric disease. The study was approved by the “Comité de Protection des Personnes Sud Méditerranée” (agreement n° 10–41). Same data were used in several previous studies (Aya Kabbara et al., 2019; Paban et al., 2019). After EEG acquisition, all participants have completed the resting-state questionnaire (ReSQ). This latter consists of 62 items organized by five main types of mental activity: visual mental imagery, inner language, somatosensory awareness, inner musical experience, and mental manipulation of numbers (Delamillieure et al., 2010). Using a scale ranging from 0 to 100%, each participant rated the percentage of time spent in each mental activity during the resting-state EEG acquisition, such that the total score for the five types of activities equaled 100%.

##### 1.2.2. Data Acquisition and Preprocessing

Each EEG session consisted in a 10-min resting period with the participant’s eyes closed (Paban et al., 2018). Participants were seated in a dimly lit room, were instructed to close their eyes, and then to simply relax until they were informed that they could open their eyes. Participants were informed that the resting period would last approximately 10 min. The eyes-closed resting EEG recordings protocol was chosen to minimize movement and sensory input effects on electrical brain activity. EEG data were collected using a 64-channel Biosemi ActiveTwo system (Biosemi Instruments, Amsterdam, The Netherlands) positioned according to the standard 10–20 system montage, one electrocardiogram, and two bilateral electro-oculogram electrodes (EOG) for horizontal movements. Nasion-inion and preauricular anatomical measurements were made to locate each individual’s vertex site. Electrode impedances were kept below 20 kOhm. The pre-processing was addressed using the same preprocessing steps as described in several previous studies dealing with EEG resting-state data (A Kabbara et al., 2018; Kabbara et al., 2017b; Rizkallah et al., 2018). Briefly, bad channels (signals that are either completely flat or contaminated by movement artifacts) were identified by visual inspection, complemented by the power spectral density. These bad channels were then recovered using an interpolation procedure implemented in Brainstorm (F Tadel et al., 2011) by using neighboring electrodes within a 5-cm radius. Epochs with voltage fluctuations between +80 μV and −80 μV were kept. Four artifact-free epochs of 40-s length were selected for each participant.

#### 1.3. Dynamic brain networks construction

For the two EEG datasets, dynamic brain networks were reconstructed using the “EEG source connectivity” method (M Hassan and Wendling, 2018) combined with a sliding window approach as detailed in (A. Kabbara et al., 2018; Kabbara et al., 2017b; Rizkallah et al., 2018). “EEG source connectivity” involves two main steps: i) solving the inverse problem in order to estimate the cortical sources and reconstruct their temporal dynamics, and ii) measuring the functional connectivity between the reconstructed time-series.

Briefly, the steps performed were the following:

1. EEGs and MRI template (ICBM152) were coregistered through the identification of anatomical landmarks by using Brainstorm (F Tadel et al., 2011).
2. A realistic head model was built using the OpenMEEG (Gramfort et al., 2010)software.
3. A Desikan-Killiany atlas-based segmentation approach was used to parcellate the cortical surface into 68 regions (Desikan et al., 2006).
4. The weighted minimum norm estimate (wMNE) algorithm was used to estimate the regional time series (Hamalainen and Ilmoniemi, 1994).
5. The reconstructed regional time series were filtered in alpha 8-13 Hz and beta 13-30Hz frequency bands, shown to be the most involved frequency bands at rest.
6. To compute the functional connectivity between the reconstructed regional time-series, we used the phase locking value (PLV) metric (Lachaux et al., 2000; Varela et al., 2001) defined by the following equation:

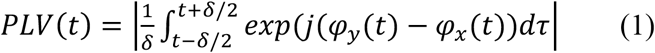

where *φ*_*y*_(*t*) and *φ*_*x*_(*t*) are the unwrapped phases of the signals *x* and *y* at time *t.* The Hilbert transform was used to compute the instantaneous phase of each signal. *δ* denotes the size of the window in which PLV is calculated. Dynamic functional connectivity matrices were computed for each epoch using a sliding window technique (A. Kabbara et al., 2018; Kabbara et al., 2017b; Rizkallah et al., 2018). It consists in moving a time window of certain duration *δ* along the time dimension of the epoch, and then PLV is calculated within each window. As recommended by (Lachaux et al., 2000), the number of cycles should be sufficient to estimate PLV in a compromise between a good temporal resolution and a good accuracy. The smallest number of cycles recommended equals to 6. For instance, in the alpha band, we chose the smallest window length of 571 ms that is equal to 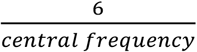.
7. To ensure equal network density for all the dynamic networks computed across time, a proportional (density-based) threshold was applied in a way to keep the top 15% of connectivity values in each network.

### 2. MEG dataset (HCP)

#### 2.1. Participants

As part of the HCP MEG2 release (Larson-Prior et al., 2013; Van Essen et al., 2012), resting-state MEG recordings were collected from 61 healthy subjects (38 women). The release included 67 subjects, but six subjects were omitted from the analysis as their recordings failed to pass the quality control checks (including tests for excessive SQUID jumps, sensible power spectra, correlations between sensors, and availability of sufficient good quality recording channels). All subjects are young (22–35 years of age) and healthy.

#### 2.2. MEG recordings and pre-processing

The acquisition was performed using a whole-head Magnes 3600 scanner (4D Neuroimaging, San Diego, CA, USA). Resting state measurements were taken in three consecutive sessions of 6 min each. Data were provided pre-processed, after passing through a pipeline that removed artefactual segments, identified faulty recording channels, and regressed out artefacts which appear as independent components in an ICA decomposition with clear artefactual temporal signatures (such as eye blinks or cardiac interference).

#### 2.3. Dynamic brain networks construction

Here, we adopted the same pipeline used by the previous studies dealing with the same dataset (Colclough et al., 2015). Thus, to solve the inverse problem, we have applied a linearly constrained minimum variance beamformer (Van Veen et al., 1997). Pre-computed single-shell source models are provided by the HCP and the data covariance were computed separately in the 1–30 Hz and 30–48 Hz bands as in (Colclough et al., 2016a). Data were beamformed onto a 6 mm grid using normalized lead fields. Then, source estimates were normalized by the power of the projected sensor noise. Source space data were filtered in alpha (8-13Hz) and beta bands (13-30Hz). After obtaining the regional time series on the basis of the Automated Anatomical labelling atlas (AAL)(Tzourio-Mazoyer et al., 2002), a symmetric orthogonalization procedure (Colclough et al., 2015)was performed for signal leakage removal. To ultimately estimate the functional connectivity between regional time series, we used the amplitude envelope correlation measure (AEC) (Brookes et al., 2012). This method briefly consists of 1) computing the power envelopes as the magnitude of the signal, using the Hilbert transform, and 2) measuring the linear amplitude correlation between the logarithms of ROI power envelopes. Finally, a sliding window (length = 6 sec, step = 0.5 sec) was applied to construct the dynamic connectivity matrices. This sliding window has been previously used to reconstruct the dynamic networks derived from MEG data(O’Neill et al., 2017). Also, matrices were thresholded by keeping the strongest 15% connections of each network.

### 3. Extracting modular brain states

Modularity refers to the extent to which a network can be separated into modules or communities highly intra-connected and weakly inter-connected (Sporns and Betzel, 2016). To track the transient changes of the brain modular networks over time, we used our recent proposed algorithm (A. Kabbara et al., 2019) that aims to extract the main modular structures (i.e modular states) that fluctuate repetitively across time. Each modular state reflects unique spatial modular organization. Briefly, the algorithm consists of applying the following steps:

- Decompose each temporal network into modules using the consensus modularity approach, which aims to address the degeneracy problem (the fact that different modularity algorithms produce different results when repeatedly applied on the same network) (Bassett et al., 2013; Kabbara et al., 2017b). The consensus modularity algorithm produces a set of partitions acquired from the Newman algorithm (Girvan and Newman, 2002)and Louvain algorithm repeated for 200 runs (Blondel et al., 2008).Then an association matrix of N × N (where N is the number of nodes) is computed by counting the number of times two nodes are assigned to the same module across all runs and algorithms. The association matrix is then compared to a null model association matrix computed from a permutation of the original partitions, and only the significant values are retained (Bassett et al., 2013). To ultimately obtain consensus communities, we re-clustered the association matrix using Louvain algorithm.
- Assess the similarity between the temporal modular structures using the z-score of Rand coefficient, a value between 0 (totally different structures) and 1 (identical structures) as proposed by (Traud et al., 2008). This step generated a T × T similarity matrix where T is the number of time windows.
- Cluster the similarity matrix into modular states (MS) using the consensus modularity method. This step associates common temporal modular structures into the same community. Hence, the association matrix of each MS is computed using the modular affiliations of its corresponding networks.

### 4. Extract the dominant modules

It is noteworthy to mention that the same module may be present in several distinct MSs. In the present study, we aim to examine the degree of resemblance between each unique module and the RSNs found in literature. Thus, we added to the algorithm two automatic steps:

1. Detect the common modules between all modules derived from MSs. We consider that a module is the same if more than 80% of its nodes are reserved. These extracted modules are considered as the representative “dominant communities”.
2. Associate each dominant module with one or several RSNs. To do that, we formed different masks or templates; where each mask is related to a RSN or combination between different RSNs. Then, an overall match for each module with each template is calculated. If the overlap between the module and a template is higher than 80%, the module is ultimately associated to the considered template.

Adding these steps has two major benefits i) it provides the dominant communities for each subject and ii) it standardizes the extracted communities as “prototype networks (i.e RSNs)” which allows analyzing the consistency of the derived modules at the group-level and validates the single subject results.

### 5. Quantification

For each dominant module, two metrics were computed:

1. The temporal fractional occupancy (FO) which represents the total time spent by each module as measured by percentage. Thus, a high value of FO reflects high temporal dominance of the module.
2. The mean dwell time (DT) defined as the average number of consecutive windows spent in a specific module. A module with a high DT is thus considered as a stable or “steady” module, compared to modules with low DT that are considered as “transient” modules.

### 6. Statistical tests

In order to investigate whether the observed brain modules are related to subjective internal thoughts and feelings experienced during resting-state acquisition, we have assessed the statistical relationships between the occurrence of modules and the phenotypes of cognition measured by the Resting-State Questionnaire (rsQ). More specifically, Pearson’s correlation between the five main indices derived from the rsQ (i.e visual mental imagery, inner language, somatosensory awareness, inner musical experience, and mental manipulation of numbers) and the fractional occupancies of the derived modules were computed for the 57 participants provided by Dataset 2. To consider the multiple comparisons problem (between the five types of mental activity, and the 11 modules), *p*-values were corrected using Bonferroni procedure (Bland and Altman, 1995) yielding an adjusted threshold of *p* < 0.0009.

### 7. Code availability

Data pre-processing was done using automagic Matlab toolbox https://github.com/methlabUZH/automagic (Pedroni et al., 2019) for dataset 1. Brainstorm toolbox (Franois Tadel et al., 2011) was used to pre-process the signals of dataset 2, and to reconstruct the regional time series using wMNE. To estimate the head model, OpenMEEG (Gramfort et al., 2010) software was used. Brain Networks estimation of EEG data (datasets 1,2) was done using Matlab. Beamforming construction and networks estimation of MEG data (dataset 3) was performed using the megconnectome pipeline package https://www.humanconnectome.org/software/hcp-meg-pipelines. The Matlab code developed to extract the modular brain states is publicly available at https://github.com/librteam/Modularity_algorithm_NN. BrainNetViewer (BNV) (Xia et al., 2013) https://www.nitrc.org/projects/bnv/ was used for networks visualization. Other homemade codes were also developed for statistical tests, and quantitative evaluation.

### 8. Data availability

The data used here are all available. The dataset 1 can be found on http://fcon_1000.projects.nitrc.org/indi/cmi_healthy_brain_network/sharing_neuro.html, the dataset 2 can be available upon a simple request to the correspondent author and the dataset 3 is available on https://db.humanconnectome.org/.

## Results

We conducted our analysis on three independent datasets: 1) Resting state EEG data provided from the Healthy brain network biobank comprising 444 subjects, 2) Resting state EEG data acquired from 56 subjects and 3) Resting state MEG data provided from the Human Connectome Project including 61 subjects. The dynamic functional connectivity networks were assessed for each subject using EEG/MEG source connectivity method (Mahmoud Hassan and Wendling, 2018b). For EEG datasets (datasets 1 and 2), we used the weighted Minimum Norm Estimate (wMNE) followed by phase-couplings as recommended by previous EEG studies (Hassan et al., 2016, 2014). For MEG dataset (dataset 3), we used the beamforming approach followed by envelope-couplings, as recommended in previous MEG studies(O’Neill et al., 2017; Tijms et al., 2013), with correction for spatial leakage in order to reduce the effects of volume conduction. Afterwards, a sliding window technique was applied to form a continuous series of snapshots characterizing the evolution of each individual’s functional brain network (see Materials and Methods for details about the construction of the functional networks). Then, we applied the modularity-based algorithm that takes as input the tensor of dynamic networks and produce the modular states (MSs) that fluctuates over time, where each MS represents a unique modular topology. Briefly, the algorithm finds the modular structures sharing the same topology by quantifying the similarity between all the computed temporal partitions. We then identify the individual dominant modules (DMs), that is, the common modules between the derived MSs. Finally, each dominant module was associated to one or several RSNs. The full pipeline of the study is illustrated in Figure 1.

### 15 states were identified for the first database

Figure 2 illustrates the 15 dominant modules derived from all the 444 subjects in alpha band. It also reports the percentage of subjects exhibiting each of the modules. One can realize that three derived modules are related to the DMN: POST-DMN including the posterior components of the DM system (posterior cingulate, parahippocampal, precunues, and inferior parietal lobule regions), ANT-DMN including the anterior components of DMN (prefrontal regions) and DMN which represents the large module integrating both posterior and anterior parts into the same module. Also, three temporal modular configurations are depicted: L-TEMP, R-TEMP representing the left and right superior and inferior temporal regions, respectively, and TEMP that combines both left and right temporal modules. Overall, the modules ordered from the highest to the lowest percentage of subjects are: DMN (present in 93% of subjects), POST-DMN (present in 88% of subjects), VIS-visual network (present in 84% of subjects), ANT-DMN (present in 85% of subjects), SMN-somatomotor network (present in 82% of subjects), DAN-dorsal attentional network (present in 78% of subjects), L-TEMP (present in 72% of subjects), TEMP (present in 58% of subjects), SAN-salience network (present in58% of subjects), AUD+VIS- a module that combines both auditory and visual networks (present in 58% of subjects), DMN+FPN – a module that combines both the default mode and the frontoparietal networks (present in 58% of subjects), DAN+VIS- a module that combines DAN and VIS (present in 43% of subjects), R-TEMP (present in 157 among 444 subjects), FPN (present in 35% of subjects), AUD+VIS+DAN- a module combining AUD, VIS and DAN networks (present in 27% of subjects).

**Figure 2.**
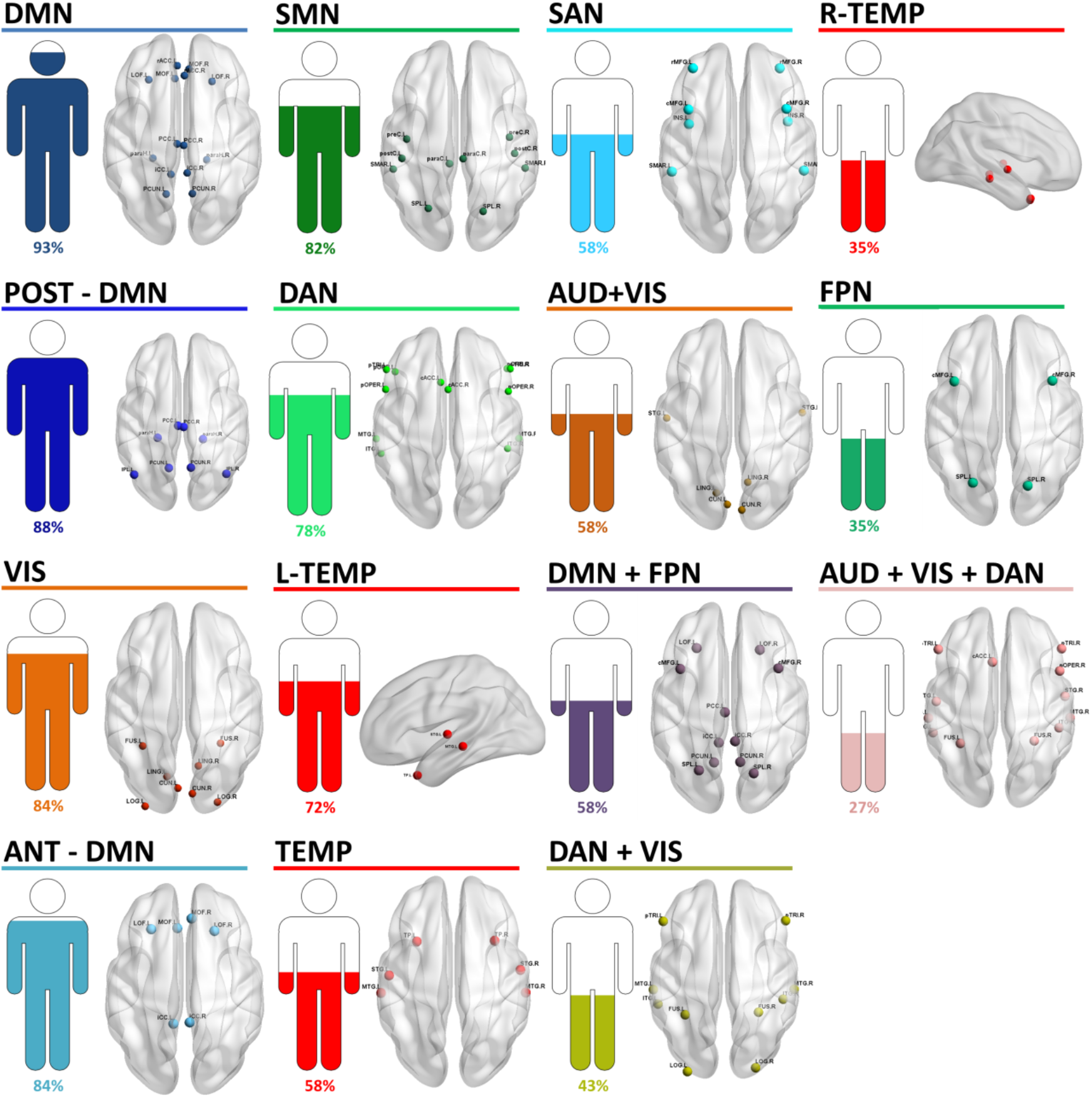
Results of Dataset 1 obtained in alpha band: The derived dominant modules and their corresponding percentage of subjects.

### 11 states were identified for the second database

According to the second dataset (Figure 3), 11 modules are extracted from the 57 subjects in alpha frequency band. These modules ordered by the percentage of subjects are: POST-DMN (present in 94% of subjects), VIS (present in 91% of subjects), DAN (present in 87% of subjects), DMN (present in 84% of subjects), L-TEMP (present in 82% of subjects), ANT-DMN (present in 81% of subjects), SMN (present in 68% of subjects), AUD+VIS (present in 68% of subjects), DAN+VIS (present in 50% of subjects), TEMP (present in 45% of subjects), DMN+CCN a module that combines DMN with cognitive control components (present in 21% of subjects). Results were also consistent among several threshold values of the functional connectivity matrices (see Table S1) and at beta frequency band (see Table S3).

**Figure 3.**
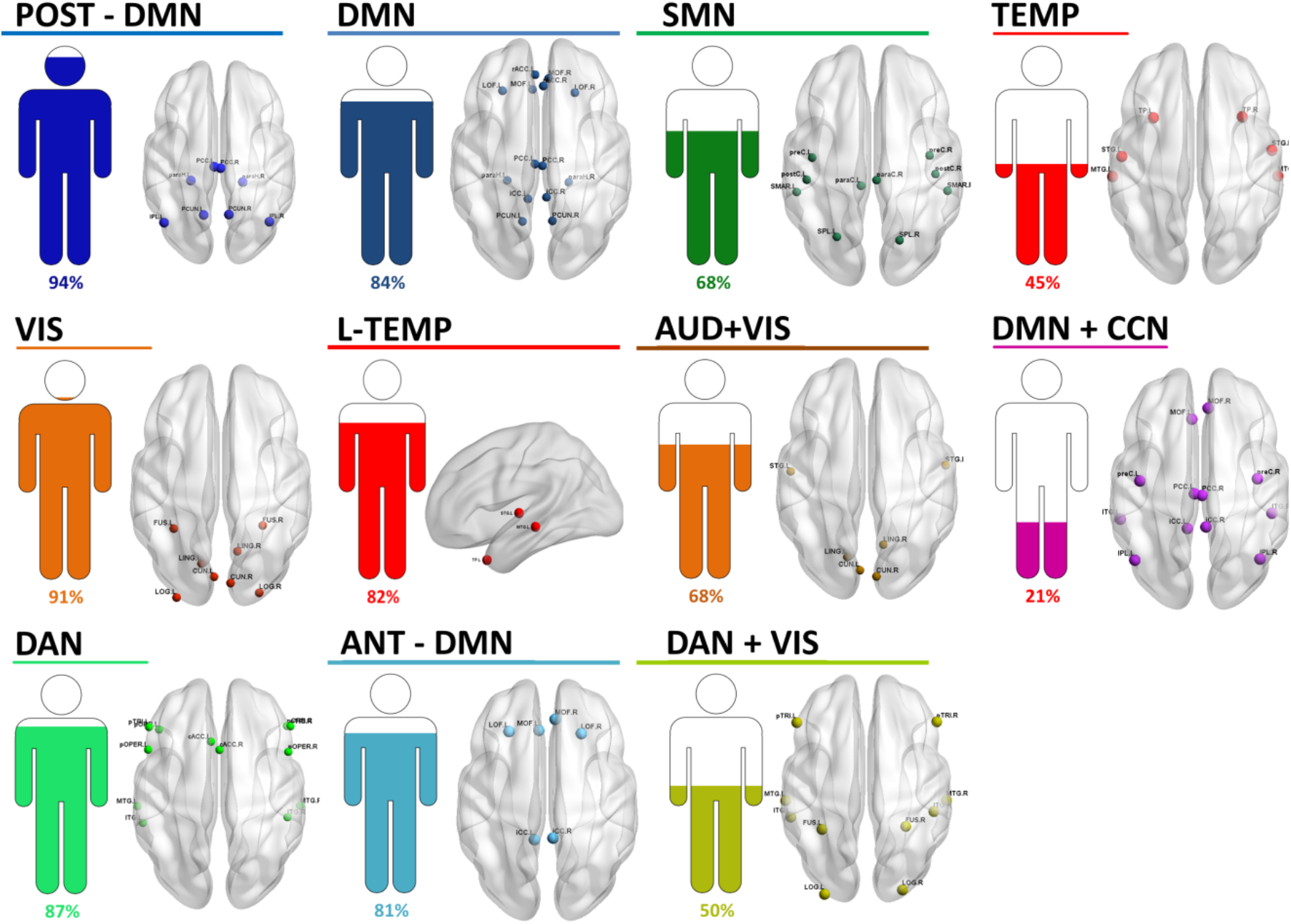
Results of Dataset 2 obtained in alpha band: The derived dominant modules and their corresponding percentage of subjects

### 9 states were identified for the third database

In figure 4, we illustrate the results obtained in alpha band for the third dataset showing 9 modules derived from the 61 subjects treated, and ordered by the percentage of subjects as follows: DMN (present in 100% of subjects), POST-DMN (present in 92% of subjects), VIS (present in 85% of subjects), L-TEMP (present in 78% of subjects), SAN (present in 72% of subjects), DAN (present in 71% of subjects), SMN (present in 56% of subjects), AUD+VIS (present in 49% of subjects), DAN+VIS (present in 43% of subjects). Results were consistent among several threshold values of functional connectivity matrices (see Table S2) and at beta frequency band (see Table S4).

**Figure 4.**
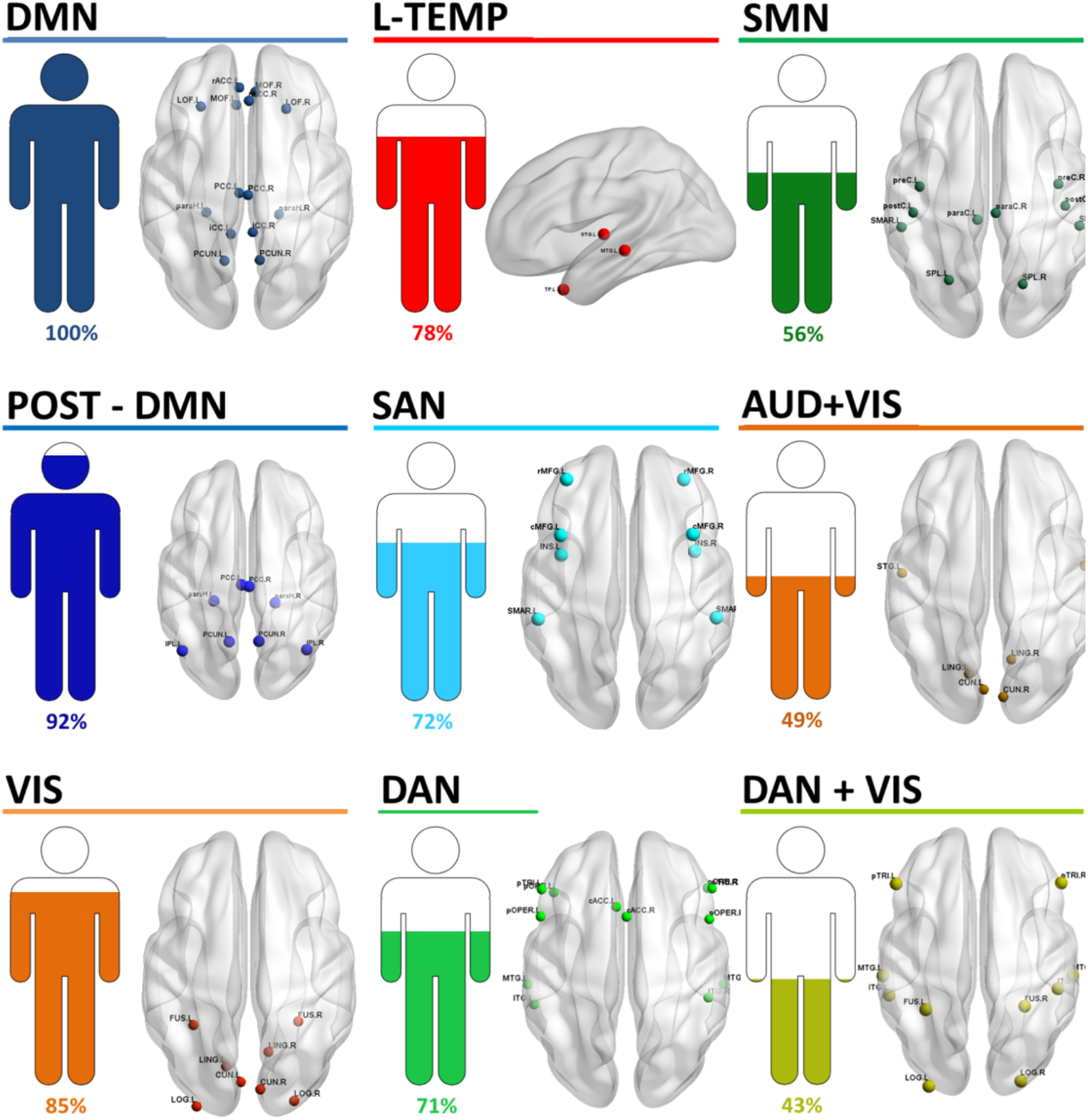
Results of Dataset 3 obtained in alpha band: The derived dominant modules and their corresponding percentage of subjects

In summary, the results obtained for the three datasets revealed fluctuating modules concordant with the well-known RSNs. In particular, one can notice that the default mode network is the most consistent network among subjects in all datasets (reflected by the highest percentage of presence over subjects). Results also showed that some RSNs present various modular topologies over time such as DMN and temporal networks. In addition, modules that combine several RSNs are observed during time, reflecting cross-network interactions, see discussion for more details.

### Dwell time and the fractional occupancy

In order to quantify the temporal characteristics of each module, two metrics were computed: dwell time (DT), i.e. the amount of average number of consecutive windows spent in a module and the fractional occupancy (FO) reflecting the proportion of time spent in each module. Figure 5 shows an example of the results of a typical subject selected from the first database. Figure 5.A shows the occurrence of ANT-DMN (FO = 79%, DT = 23%) where most of the regions are associated to the anterior part of DMN. The second detected module, referred as DMN (FO = 11%, DT = 3%), shows integration between the orbito-frontal, the anterior cingulate regions, the posterior cingulate and the precunues regions. Interestingly, a module affiliated to POST-DMN (FO = 7%, DT = 3%) incorporates the posterior cingulate, the precunues and the parahippocampal regions. In addition, SMN (FO= 48%, DT = 10%) was illustrated in the fourth module revealing the post-central, the pre-central, the paracentral, the supramarginal and the superior parietal lobule. The anterior cingulate regions were also present in this module. The fifth module showing the interactions between the lateral occipital regions, the cuneus and the lingual regions is clearly associated to VIS network (FO = 59%, DT = 12%). One can also notice the presence of the DAN (FO = 23%, DT = 7%) combining the anterior cingulate, the paratriangularis, the paraorbitalis, the middle temporal gyrus, the transverse temporal, and the pericalcarine regions. Figure 5.B illustrate the temporal fluctuation of the different modules across time.

**Figure 5.**
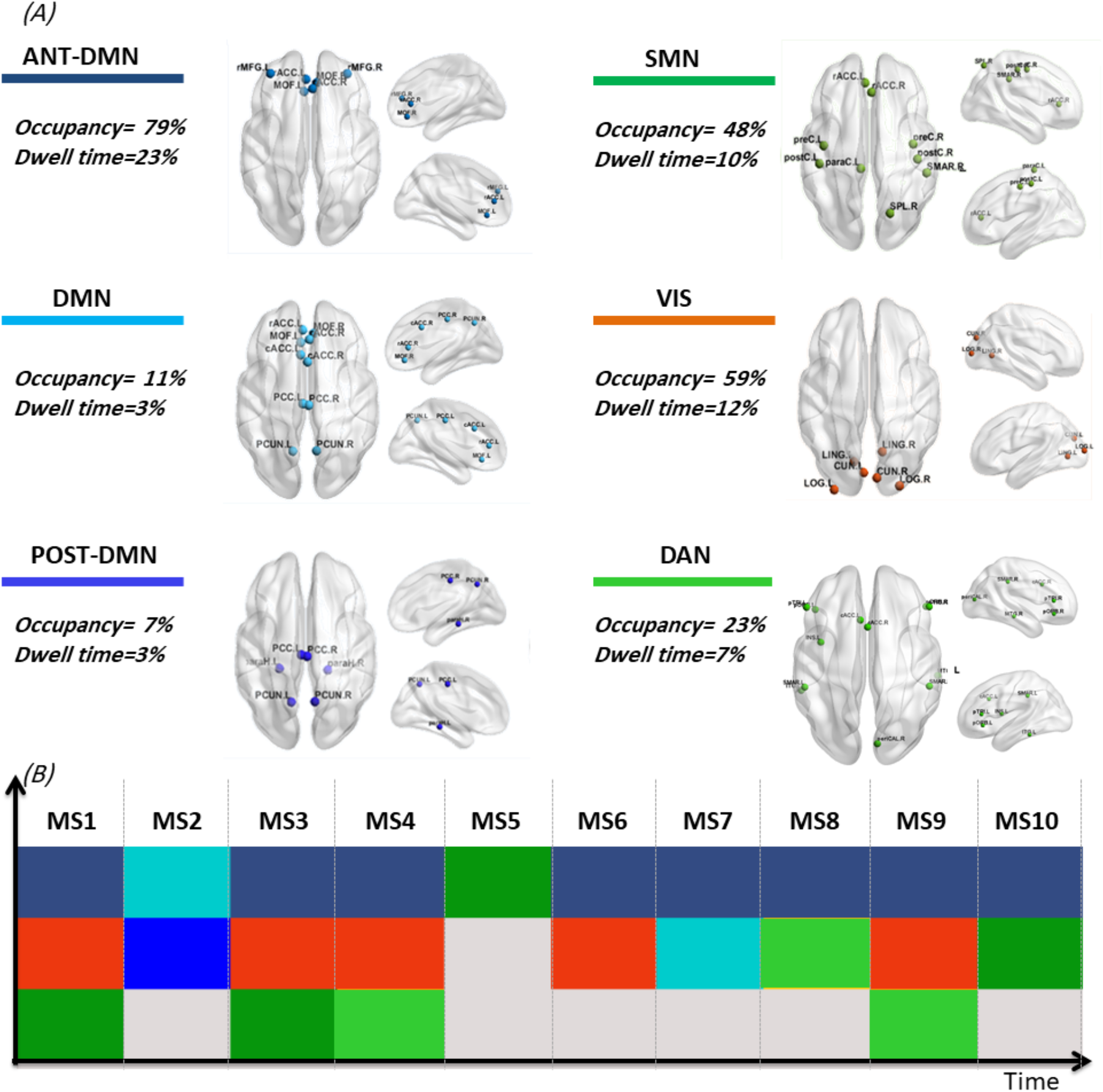
Results of a typical subject. (A) The spatial representation of the dominant modules. (B) The temporal fluctuations of the derived modules.

Figure 6 reports the fractional occupancies and the mean dwell durations of the dominant modules obtained for each dataset. One can clearly realize that DMN (or one of its modular configurations) has the highest TO and DT over all datasets. The VIS network is also revealed significant in terms of FO in datasets 2 and 3. According to the DT, SMN and SAN are depicted as significantly stable modules in dataset 1, while VIS is considered as significant in dataset 2.

**Figure 6.**
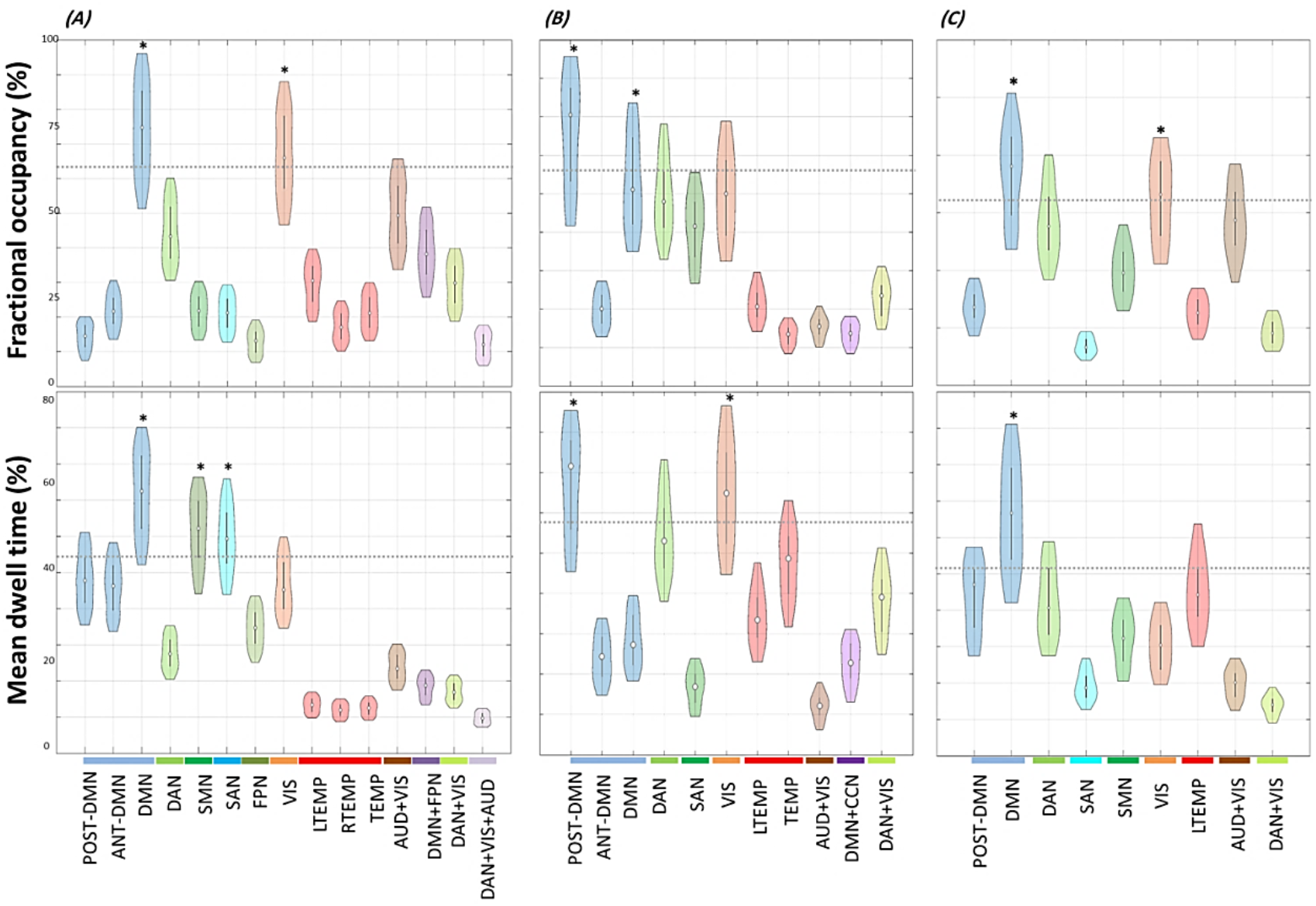
The violin plots showing the fractional occupancy and the mean dwell time of the derived modules obtained for (A) Dataset1, (B) Dataset2 and (C) Dataset3. The horizontal dashed line that appears in each plot denotes the mean plus two standard deviations. * mark the significant modules (average > mean value + 2 standard deviations).

In summary, the results obtained from all datasets points at stability of DMN and its role as a functional core network during rest, see discussion section for more details.

### Correlation between the derived modules and resting-state cognition

Finally, we seek at understanding if there is any correlation between the derived modules and the subject internal thoughts experienced during resting-state acquisition measured by the Resting-State Questionnaire (rsQ). Only such data was available for dataset 2. More specifically, the five main indices derived from the rsQ (i.e visual mental imagery, inner language, somatosensory awareness, inner musical experience, and mental manipulation of numbers) were correlated with the FO of each module.

Figure 7 reports significant positive correlations between the visual mental imagery and the fractional occupancy of VIS (*p*_*Bonferroni corrected*_ = 0.0006; *R* = 0.44), AUD (*p*_*Bonferroni corrected*_ = 0.0008; *R* = 0.451) and DAN (*p*_*Bonferroni corrected*_ = 0.0001; *R* = 0.48) obtained in alpha band. In beta band, results show positive correlations between the visual imagery and the FO of AUD and DAN (see Figure S2). Also results were consistent across other thresholds (see Figure S1).

**Figure 7.**
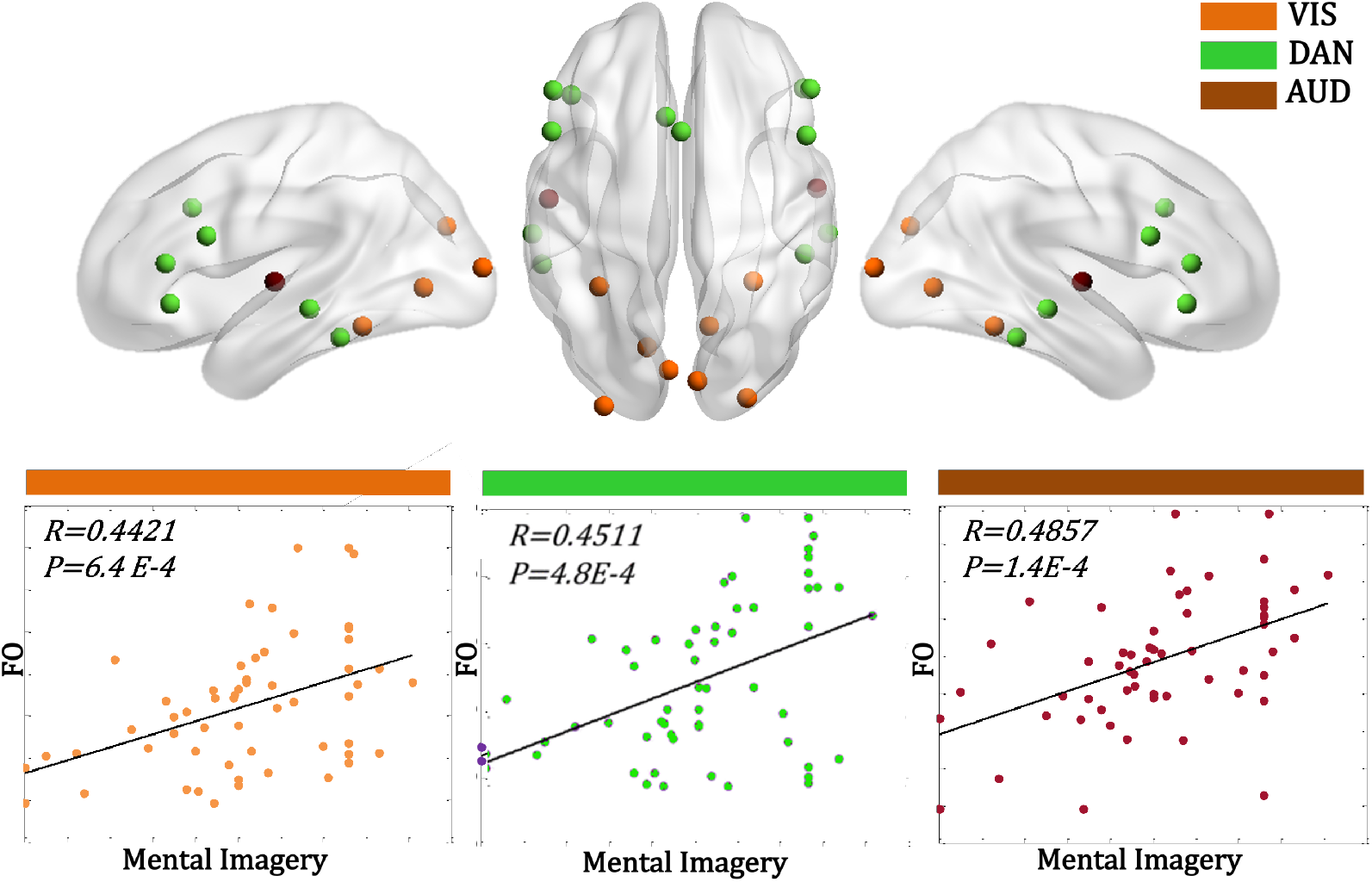
Significant correlations (Bonferroni corrected) between visual mental imagery and the fractional occupancies (FOs) of VIS, AUD and DAN modules in alpha band (dataset 2).

In summary, these results showed that individual variability in the visual imagery experienced during acquisition was positively related to the occupancy of specific modules, mainly VIS, DAN and AUD.

## Discussion

We show how fast changes in the modular architecture shape the spontaneous brain activity. Here, we used a recently developed algorithm that extracts the repetitive modular brain states alternating over time. Unlike the traditional approaches, the distinctive feature of the applied method resides in tracking the modular variations of brain networks. The method was applied on three independent EEG/MEG datasets and showed that the well-known RSNs show continuous modular changes reflected by a process of separation and merging within and between the resting networks.

In particular, DMN switches dynamically its modular topology, in line with many studies suggesting that DMN is decomposed into subcomponents, mainly anterior and posterior portions (Andrews-Hanna et al., 2007; Moussa et al., 2011; Wens et al., 2019). Also, the process of association and dissociation within DMN components is also revealed by Allen et al.(Damaraju et al., 2014) where brain states were described using K-means clustering. More importantly, several studies also showed that the dynamic states transition leads to the inclusion of some FPN regions in the DMN in some brain states(Damaraju et al., 2014; Liu et al., 2019), which was also obtained in our study (results of dataset 1). Similarly, the temporal network alternates its reconfiguration between left, right and complete modules. This finding is in concordance with previous results depicting the left part of the temporal network as an independent network state (Baker et al., 2014). The dynamic modular behavior of the resting brain was also revealed by the occurrence of modules integrating different RSNs. For instance, DAN expands dynamically its network to include the visual components. The dynamic inclusion of these networks echoes the presence of high correlation between them, which was supported in previous studies (Damaraju et al., 2014; Liu et al., 2019).

Our findings agree with previous studies suggesting that some brain regions are transmodal, meaning that they are connected to different resting state networks over time (Zalesky et al., 2014). These regions are highly dynamic and change their affiliated module during time. A similar observation was obtained in a previous study where a defined hub was shown to dynamically alternate its role between provincial and connector hub (Kabbara et al., 2017b). This dynamic process of splitting and merging the different sub-systems during time allows the brain to balance segregated and integrated neural dynamics.

The DMN (in its different configurations) is the most consistent module obtained across subjects, as it shows the highest percentage of presence over subjects/datasets, with the high fractional occupancy values. These findings highlight the key important role of DMN in integrating information in spontaneous functional brain activity, in accordance with the presence of a large proportion of hubs associated to this network (de Pasquale et al., 2015; de Reus and van den Heuvel, 2013; Kabbara et al., 2017b; van den Heuvel and Sporns, 2013). Furthermore, one can realize that results from all datasets show that one of the DMN configurations holds the greatest stability over time (reflected by the highest mean dwell duration). Such findings support previous studies showing that functional connections within the rich-club core (where most regions are affiliated to DMN) exhibit the greatest stability over time. In these studies, the high temporal stability of DMN has been associated with high dependency on the underlying structural brain topology, as high similarity was obtained between resting-state functional and structural networks when the sample duration increases. In contrast to this, modules with low dwell time are transient modules exhibiting greatest variability during time and reflecting the dynamic functional coordination. In our study, the derived transient modules depend on each subject and database. Mostly, they belong to the high level cognitive networks, attentional networks and the sensory networks. In all databases, these transient modules showing significant low dwell time are those integrating many RSNs (DAN+VIS, AUD+VIS…). This can be explained as individuals dynamically engage in several mental thoughts during resting periods while imagery and mind-wandering remain the predominant activities (Delamillieure et al., 2010; Doucet et al., 2012). Thus, looking at the brain as a dynamic system where stable activity is intertwined by transient functional variabilities was supported by many studies (Honey et al., 2007; Liu et al., 2019; Van De Ville et al., 2010).

In addition, the study highlights the significant presence of the visual network showing high occupancy rate during time (results of datasets 1 and 2). Such observation can be associated with the dominance of the visual imagery activity exhibited by most subjects during resting state acquisition(Delamillieure et al., 2010). More interestingly, the individual variability in the visual imagery experienced during acquisition was revealed to be positively related to fractional occupancy of VIS (Figure 7). Similar correlations were reported by previous studies (Pipinis et al., 2017; Stoffers et al., 2015). In addition, the significant relationships assessed between the mental imagery with AUD, DAN and VIS may explain the cross-interactions observed between these networks forming one large module during time (datasets 1,2,3).

Across the three datasets, results show 9 common RSNs: DMN, POST-DMN, VIS, SMN, SAN, LTEMP, VIS+AUD, DAN+VIS. These striking consistent results are obtained independently from the technique used to record signals (EEG or MEG), the preprocessing steps (automatic in dataset 1 vs. manual in dataset 2 and 3), the source reconstruction (wMNE vs. Beamforming), the adjacency matrices thresholding value, the EEG/MEG frequency bands (alpha vs. beta), the atlas parcellation (68 Desikan Killiany vs 78 AAL) and the functional connectivity measures (phase vs. envelope couplings) as well as either with or without correcting the zero-lag correlations. This prompts us to be somewhat confident of the results we obtained and that they don’t rely on specific methodological considerations.

However, other dominant modules arise from each dataset (Figures 2,3,4). The inter-subject variability was also revealed by the percentage of subjects showing each dominant module. Among the same dataset, these individual differences are thought to be associated with variability in important cognitive and behavioral functions. This has been supported by different studies showing that the dynamic network characteristics significantly correlate with intelligence, creativity and executive function (Bassett et al., 2015; Kenett et al., 2020; Simony et al., 2016; Tompson et al., 2018). Here, between-subjects variation in the temporal characteristics of specific modules, mainly VIS, AUD and DAN, was associated with self-report rating of mental visual imagery as measured by the resting-state questionnaire. The dependence of observed brain activity on the inner thoughts and feeling experienced during resting acquisition was emphasized by multiple studies (Diaz et al., 2016; Pipinis et al., 2017; Stoffers et al., 2015). The discrepancy of results obtained from different datasets may be due to some difference in the datasets such as the sample size and age of subjects. In addition, MEG/EEG differences proved to be particularly arisen when investigating the transient resting-state functional connectivity patterns (Coquelet et al., 2020).

The relevance of alpha to beta frequency range (8–30 Hz) in driving spontaneous large-scale neuronal interactions was revealed by multiple EEG/MEG studies (Brookes et al., 2011; Hipp et al., 2012a; Kabbara et al., 2017a; Liu et al., 2010; Pasquale et al., 2010). As correlation patterns depend on the underlying oscillation frequency (Brookes et al., 2011; Hipp et al., 2012b; Siegel et al., 2012; Vidaurre et al., 2018), we have verified the reproducibility of the obtained results in these two frequency bands. The main conclusions of the study remain intact (see Table S3, Table S4, Figure S2): i) distinct modules concordant with the well-known RSNs fluctuate during time ii) the default mode network is detected as the most consistent, dominant and stable module which dynamically alternates its modular topology, iii) modules that combine several RSNs are observed during time, reflecting cross-network interactions such as DAN-VIS and iv) significant positive correlation was revealed between the fractional occurrences of some specific modules and the mental imagery. Nevertheless, slight differences were observed in the derived modules and their temporal characteristics between the two frequency bands.

From methodological viewpoint, we have adopted in each dataset the pipeline (from data processing to networks construction) used by the previous studies dealing with the same datasets. Thus, for EEG datasets, we used the wMNE/PLV combination to reconstruct the dynamic networks, as it is supported by several studies on resting EEG and two comparative studies (Hassan et al., 2016, 2014). For the MEG dataset, beamforming construction combined with amplitude correlation (and orthogonalization) between band-limited power envelops was used by multiple studies using the MEG HCP data (Brookes et al., 2012; Colclough et al., 2015; George C. O’Neill et al., 2016). The suitable window width is a crucial issue in constructing the dynamic functional networks. On the one hand, short windows do not contain sufficient information to accurately estimate connectivity. On the other hand, large windows may fail to capture the temporal changes of the brain networks. Hence, the ideal is to choose the shortest window that guarantees a sufficient number of data points over which the connectivity is calculated. This depends on the frequency band of interest that affects the degree of freedom in time series. It is also depending on the correlation measure used. In EEG datasets, we adapted the recommendation of Lachaux et al. (Lachaux et al., 2000) in selecting the smallest appropriate window length offering 6 number of ‘cycles’ at the given frequency band. The reproducibility of resting state results whilst changing the size of the sliding window was validated in a previous study (Kabbara et al., 2017b). In MEG, we used the same correlation method with the corresponding sliding window size (0.5 sec) applied in previous studies dealing with the same dataset (Colclough et al., 2016b; George C. O’Neill et al., 2016).

In this study, we used a proportional threshold (highest 15% of the edge’s weights) to remove weak connections. The stability of network-based features across proportional thresholds was supported by (Garrison et al., 2015) in contrary to absolute thresholds. In addition, applying a proportional threshold is important to ensure equal density between network derived from different time windows and subjects. Nevertheless, and in order to ensure that the obtained results are not sensitive to the threshold value, we performed our analysis across three proportional thresholds: 5%, 15%, 30%. High degree of agreement among the obtained results was found, see Supplementary Materials (Table S1, Table S2, Figure S1).

To extract the fast transient modules, we have applied the modularity-based algorithm that extracts the main modular brain states fluctuating during time(A. Kabbara et al., 2019). Other strategies aiming at identifying the connectivity states exist such as K-means clustering, ICA and PCA for isntance. However, in these frameworks, states are identified without considering the modular organization of the networks. Distinguishingly, the applied algorithm proceeds its segmentation from looking at the brain as a dynamic modular network. In a previous study, a quantitative comparison using simulated data was performed between the modularity-based algorithm, K-means clustering(Damaraju et al., 2014), independent component analysis(George C O’Neill et al., 2016) and the consensus clustering (Rasero et al., 2017). The framework used here outperformed the other techniques in terms of spatial and temporal accuracy compared.

## Acknowledgments

This work was financed by the Rennes University, the Institute of Clinical Neuroscience of Rennes (project named EEGCog). The study was also funded by the National Council for Scientific Research (CNRS) in Lebanon. Authors would also like to thank the Lebanese Association for Scientific Research (LASER) for its support.

